# Arbuscular mycorrhizae reduce stress-induced plasticity of plant functional traits. A meta-analysis study

**DOI:** 10.1101/2022.05.15.491379

**Authors:** Florencia Globbo, María José Corriale, Ayelén Gázquez, César Daniel Bordenave, David Bilenca, Ana Menéndez

## Abstract

We aimed at exploring the plant functional traits whose stress-induced plasticity is altered by the presence of AM fungi, considering the direction of their changes. We also sought for a coordinated variation of plant biomass and functional traits, during plant adaptation to environmental stressors, and the role of AM status on the variation. We performed a meta-analysis across 114 articles spanning 110 plant species or cultivars. We quantified the size effect of AM symbiosis on the stress-induced plasticity of several reported and calculated functional traits, and using linear mixed model analysis (LMM). Correlation between traits plasticity and total biomass variation were also performed through LMM. The literature search and further selection yielded seven functional traits, extracted from 114 laboratory studies, including 888 observations and 110 plant species/cultivars. Evidence for significant effects of predictor variables (type of stress, AM symbiosis and/or their interaction) on plasticity were found for three of these functional traits: leaf-area ratio (LAR), root mass fraction (RMF) and root-shoot (R:S) ratio. Our results provided evidence to accept the hypothesis that AM fungal inoculation may reduce the phenotypic plasticity of important plant functional traits leaf area ratio (LAR), root mass fraction (RMF) and root-shoot (R:S) ratio, by decreasing its magnitude. We also found a weak correlation between traits plasticity and total biomass variation. Although our literature search and data collection were intensive and our results robust, the scope of our conclusions is limited by the agronomical bias of plant species targeted by the meta-analysis. Further knowledge on non-cultivable plant species and better understanding of the mechanisms ruling resources allocation in plants would allow more generalized conclusions.

## Introduction

Soil borne abiotic stresses are considered a major source for decreases in crop yields and quality in many areas worldwide (IPCC, 2021;Yamaguchi and Blumwald, 2005, Shahbaz and Ashraf, 2013). Drought and salinity lower the soil water potential, which leads to harmful osmotic effects in plants, including slower plant growth, the impairment of some nutrients uptake and their acropetal translocation to growing plant organs (Munns, 2002; Hu and Schmidhalter, 2005). In addition to the osmotic effect, salinity may bring toxicity due to excessive Na+ accumulation and nutritional imbalances (Munns, 2005), inhibiting both cell production and cell expansion (Neumann 1995). As a result of these effects, changes in plant biomass allocation may occur.

Inversely, the symbiosis with arbuscular mycorrhizal (AM) fungi may promote plant growth by improving plant absorption of water and several important macro and micro nutrients (Al-Karaki and Al-Raddad, 1997; Liu et al., 2002; Quilambo, 2004; Chen et al., 2018). Besides, AM fungi modulate phytohormones as part of the plant’s tolerance response (Evelin et al., 2019).

The adjustment of the phenotypic expression in response to environmental stress, or to biotic interactions is known as phenotypic plasticity (Schlichting 2002; Matesanz et al., 2018). This adjustment, which is considered an attribute of the genotype, may include changes in functional traits, i.e.: the relationships between morphology and biomass allocation to different plant organs (McGill et al., 2006). Functional traits strongly influence an organism’s performance, by improving the cost-benefit in the acquisition of limiting resources, therefore alleviating the restriction effect (Freschet et al., 2015, 2018).

Major functional traits studied in plants are the leaf area ratio (LAR) and the root length ratio (RLR; Marcelis et al. 1998; Ryser 1998; Hill et al., 2006; Ostonen et al.,2007; Poorter et al., 2012; Freschet et al., 2015). LAR represents the leafiness or leaf expansion of a plant, and it is a product between the leaf mass fraction (LMF) and the specific leaf area (SLA). The RLR describes the plant potential for soil resource acquisition, and it is composed by the root mass fraction (RMF) and the specific root length (SRL). These functional traits may be affected by drought and/or salinity, according to several field and laboratory studies (Bayuelo-Jimenez et al., 2003; Meier et al., 2008; Poorter et al., 2009; Miranda et al., 2010; Rewald et al., 2013; Nguyen et al., 2014; Uchiya et al., 2016; Eziz et al., 2017). In contrast, reports about the effect of AM inoculations on SLA and LMF in plants grown under drought or saline stress are less abundant and show dispar results according to the plant species (Miranda et al., 2011; Abdel-Fattah et al., 2013; Romero-Munar 2019).

Plastic responses can change if plant perception of the resource’s limitation is modified by the increase of plant acquisition capacity for that resource (Freschet et al., 2018; Chapin et al., 1987; van Kleunen & Fischer, 2005; Valladares et al., 2007). On this basis, and as AM fungi generally improve plant nutrition and water acquisition, it could be hypothesized that AM-colonized plants would have a lower plastic response in traits that are crucial for plant adaptation to edaphic restrictions, compared with uninoculated controls. A few individual case studies support this hypothesis, such as the finding that AM fungi led to low plasticity of root-related functional traits in *Zea mays* plants confronted to a phosphorus supply gradient (Wang et al., 2020), and *Lotus tenuis* exposed to high salinity (Echeverria et al., 2008). However, comprehensive studies aimed at understanding the effects of the AM-symbiosis on the stress-induced plasticity (magnitude and direction) of functional traits have not been so far conducted.

Meta-analysis has been used to understand the response of plant functional traits to the environment (Poorter et al., 2010; 2012), and to uncover general trends in the effectiveness of AMF improvement of plant growth and ions homeostasis (Hoeksema et al., 2010; Veresoglou et al., 2012; Augé et al., 2014; Chandrasekaran et al., 2014; He et al., 2014; Jayne and Quigley, 2014; Yang et al., 2015; Chandrasekaran et al., 2016, Chaudhary et al., 2016). Here we performed a meta-analysis on experimental studies across multiple plant species, where the effects of AM-symbiosis and environmental stress on growth and/or plant functional traits have been tested. Our aim was to explore the plant functional traits whose stress-induced plasticity is altered by the presence of AM fungi, considering the direction of their changes. Here we also wondered if plant biomass and functional traits have a coordinated variation during plant adaptation to environmental stressors, and if the AM status has any effect on that variation.

## Materials and Methods

### Data source

As targets, we have searched for all articles dealing with the response of mycorrhizal versus non-mycorrhizal plants when exposed to drought or salinity environments, encompassing the period from January 1987 to May 2022. The search engines used were Scopus and Google Scholar. The search terms were “arbusc*” and “mycorrh*”, in combination with some of the following words: “drought” or “water stress”, “salinity” or “saline”. Citations and references of selected papers were also checked to ensure a comprehensive list of studies. The search rendered 1136 articles.

## Articles and data screening criteria

Retrieved articles were screened so that they met the following predefined criteria: 1-data obtained from experiments in greenhouse or microcosms, 2-minimum number of replicates=4, 3-non-mycorrhizal and non-stress control treatments must be included, 4-plant growth substrate before AM inoculation should be sterilized in order to achieve full control of AM propagules (articles using fungicides were excluded), 5-contained at least one plant growth parameter (biomass or plant height), 6-plants were none-extremophiles, 7-published in English or Spanish, 8-peer-reviewed, with full text available.

The screening rendered 114 articles fulfilling the established criteria. Treatments including AM fungi interacting with other microorganisms, or other confounding experimental treatments were discarded. AM species were reclassified according to the taxonomic scheme of Schüßler and Walker (2010) and Redecker et al. (2013). In the case of scientific articles where several types of stress, AM identity, and plant species or (cultivars) combinations were recorded, each combination was considered as an independent observation, and therefore analyzed as a separate study. In addition, when stress treatments included gradients, plasticity was calculated for every stress level in the gradient.

### Data collection

We extracted data from tables and figures. In the case of figures, the web base tool https://automeris.io/WebPlotDigitizer/ was used. We obtained mean and sample size (N) of morphometric plant growth parameters: total, root and shoot biomass, root length and leaf area. Total dry weight (TDW) was calculated when it was not provided, if possible. The following functional traits were calculated for each plant: LAR (total leaf area per TDW, cm^2^.g^-1^), SLA (leaf area per unit leaf mass, cm^2^.g^-1^), LMF (total leaf mass per TDW, g.g^-1^), RLR (total root length per TDW, m.g^-1^), SRL (total root length per root mass, m.g^-1^), RMF (root mass per TDW, g.g^-1^) and root:shoot ratio (R:S; total root mass per unit of shoot mass, g.g^-1^).

Plasticity was estimated for each functional trait, as the percentage of change in mean value from control to stress environment (Valladares et al., 2006; Molina-Montenegro and Naya, 2012; Matesanz et al., 2017), as follows:

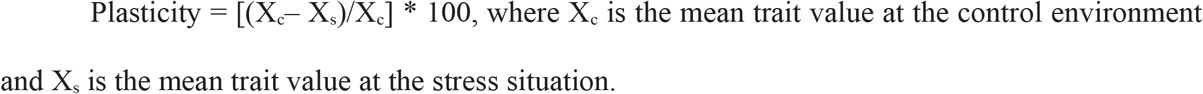

A similar calculation was used to estimate the stress-induced plant biomass variation:

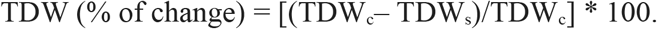

Meta-analysis of plasticity data was performed by fitting linear mixed-effects models (LMMs; Stram, 1996) in R-software version 4.0.0 (R Core Team, 2019), using the function lmer from the lme4 package (Bates et al., 2015). Assumptions of normality and homogeneity of variance were tested with Shapiro-Wilk and Levene’s tests, respectively. When necessary, we modelled heteroscedasticity with different variance structures (varIdent, varPower and varExp) fitting generalized least squares (GLS) models, in the nlme package (Pinheiro et al.,2016). In all models we included AM symbiosis, type of stress and their interaction as fixed effects. We were not interested in analyzing the effect of plant species on traits plasticity, but since they can explain part of the variability, it was incorporated to the model as a random factor, along with the scientific article, to checked for publication bias (Yang et al., 2015). We also incorporated the number of replicates used in each experiment / scientific article as an offset in the models.

In the following step, we tested for associations between the salt-induced variation in plant biomass and functional trait plasticity, in AM and non-AM plants. For this purpose, plasticity data from different stresses was pooled and fitted to multiple regression analysis using general mixed linear models function lmer from the lme4 package (Bates et al., 2015). The models included the TDW percentage of change as a dependent variable, and two independent variables, symbiosis and the percentage of change of each functional trait, in interaction. We incorporated the scientific article as a random effect and the number of replicates used in each experiment / scientific article as an offset in the models.

In all cases, model selection was carried out by a backward-stepwise elimination, using likelihood ratio tests criteria to compare hierarchically nested models, and to detect which terms should be dropped. At each step, the interaction or main effect with the highest P-value (based on χ2 Wald’s test) was identified and removed from the model if above an a priori 0.05 threshold. This process was concluded where all remaining effects had p < 0.05. Additionally, we used exhaustive model selection on the complete models (Burnham and Anderson 2002), with Akaike Information Criterion (AIC). Post hoc contrasts to assess effects and significance between fixed factors were conducted on models using the emmeans function in the emmeans package version 1.4 (Lenth 2019), with significance level of 0.05. All graphs were produced using either the base package or the ggplot2 package version 3.2.1 (Wickham 2016).

## Results

Effects of environmental stresses and arbuscular mycorrhization on phenotypic plasticity. The literature search and further selection rendered seven functional traits, extracted from 114 laboratory studies, including 888 observations and 110 plant species/cultivars (Supplementary Table 1).

Evidence for significant effects of predictor variables on plasticity were found for three of these functional traits: LAR, RMF and R:S ratio (Supplementary Table 1). Most of the variances not explained by the predictors (type of stress, AM symbiosis and their interaction) in the selected models were explained by the variability across articles, or it was just residual (Supplementary Table 2), whereas plant identity contribution to these variances was negligible.

Selected models for LAR revealed that the effect of AM symbiosis on this trait plasticity depended on the stress type (significant interactions t=2.62; p=0.01; Table 1; Fig. 1). Upon salinization, AM inoculated plants displayed a 28.6% lower magnitude of averaged LAR plasticity compared with non-AM ones, whereas no significant difference due to AM symbiosis was detected for drought experiments. Across plant species, values of LAR plasticity as response to salt-stress tended to be less negative with AM fungi treatment, being positive in most cases (Figure 1 II).

**Table 1.**
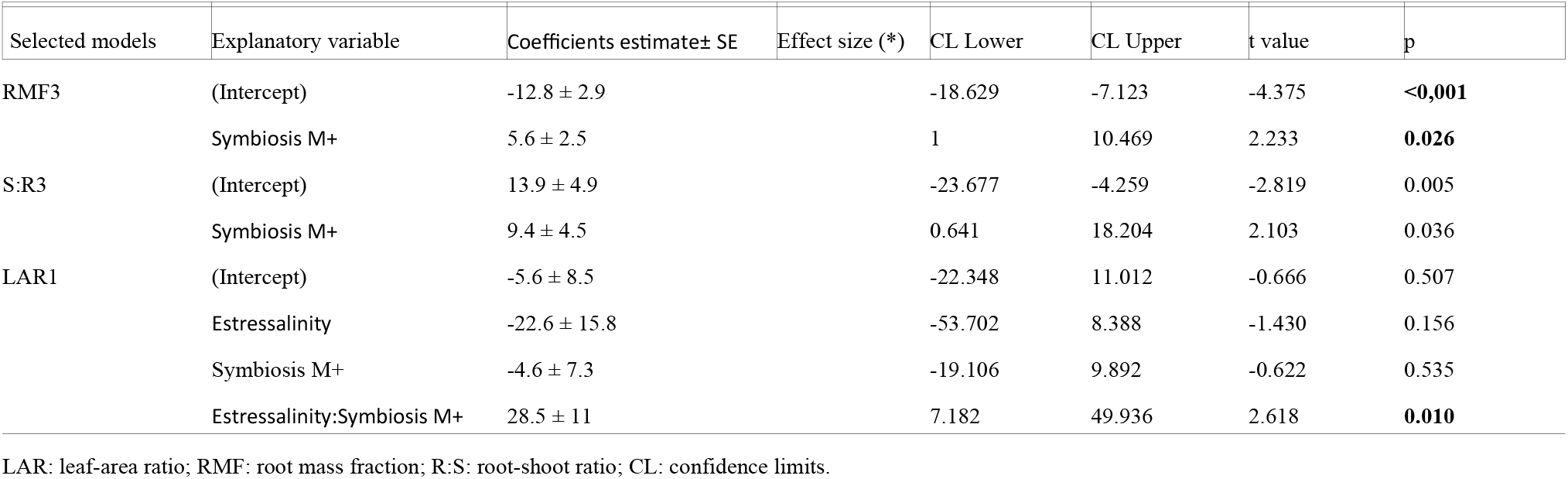
Results of the best-fit LMM testing the effects of AM symbiosis and the stress type on plant funtional traits. See Supplementary Table 1 for model comparisons.

**Figure 1.**
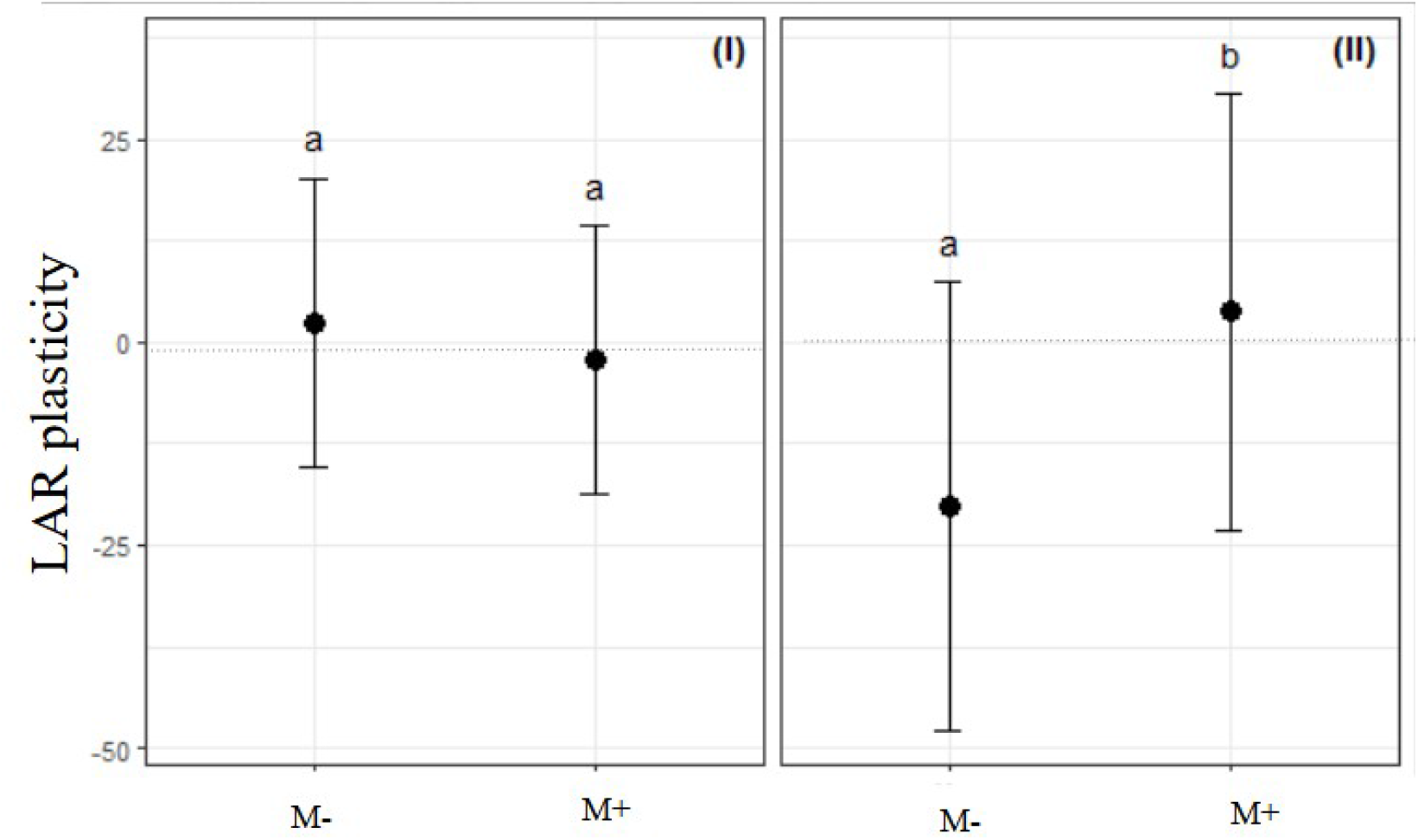
Effects of symbiosis on LAR plasticity under drought (I) and salinity (II). M-: non-mycorrhizal control. M+: mycorrhized plants.

Selected models revealed AM symbiosis as a single predictor of RMF and R:S plasticities (respectively t=2.23, p=0.026 and t=2.10, p=0.036; Table 1). Across plant species, the presence of AM fungi had a size effect of 5.6% and 9.4% reduction in the magnitudes of RMF and R:S plasticities (see coefficient estimates Table 1; Fig. 2). In the absence of AM fungi, plasticity averages of these traits were lower (more negative) compared with the AM symbiosis situation, meaning that as response to the studied stressors, non-mycorrhizal plants invested proportionally more resources in root biomass than in above ground structures.

**Figure 2.**
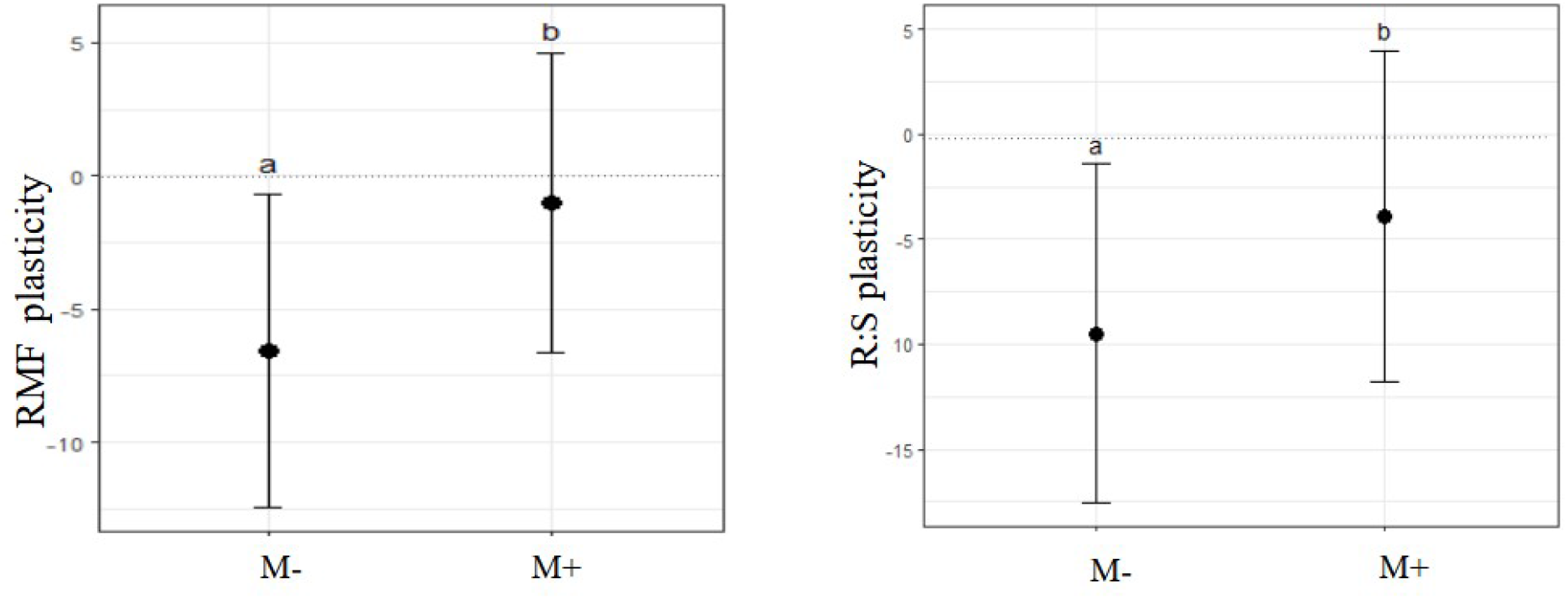
Effect of symbiosis on RMF and R:S ratio. M-: non-mycorrhizal control. M+: mycorrhized plants.

As expected, the great majority of the plant species included in the meta-analysis decreased their TDW upon the stress treatment, regardless of the presence of AM fungi (Supplementary Table 5). Evidence for significant associations between stress-induced variation of plant biomass and trait plasticity were obtained for functional traits LAR, RLR, RMF, and R:S ratio, although their regression slopes were mild (−0.313, -0.194, -0.131, -0.089 for LAR, RLR, RMF and R:S, respectively; Table 2; Supplementary Table 4). These relationships showed that higher values of stress-induced increases in these functional traits (more negative plasticity) were associated with more pronounced TDW stress-induced decreases (more positive TDW percentage of change), whereas stress-induced reductions in values of functional trait (more positive plasticities) were associated with lower TDW reductions induced by stress.

**Table 2.** Results of the best-fit LMM testing the for associations between TDW variation and plasticity of functional traits. Data from salinity and drought stresses were pooled. See Supplementary Table 4 for model comparisons.

| Selected models | Explanatory variable | Coefficients estimate± SE | CL Lower | CL Upper | t value | p |
| --- | --- | --- | --- | --- | --- | --- |
| LAR3 | (Intercept) | 10.7 ± 4.2 | 2.238 | 19.336 | 2.527 | **0.023** |
|  | LAR | *-0.313 ± 0.058* | -0.431 | -0.200 | -5.410 | **<0.001** |
| RLR1 | (Intercept) | 32.0 ± 4.7 | 22.755 | 41.099 | 6.793 | <0.001 |
|  | RLR | *-0.194 ± 0.080* | -0.350 | -0.034 | -2.423 | 0.019 |
| | Symbiosis M+ | -16.1 ± 5.6 | -27.963 | -5.086 | -2.853 | 0.006 |
| | RLR:Symbiosis M+ | -0.275 ± 0.120 | -0.519 | -0.040 | -2.294 | **0.026** |
| RMF2 | (Intercept) | 26.4 ± 2.7 | 21.001 | 31.858 | 9.561 | **<0.001** |
|  | RMF | *-0.13 ± 0.04* | -0.216 | -0.046 | -3.028 | 0.002 |
| | Symbiosis M+ | -16.1 ± 5.6 | -10.459 | -2.29 | -3.058 | 0.002 |
| R:S2 | (Intercept) | 26.2 ± 2.9 | 20.433 | 32.016 | 8.895 | **<0.001** |
|  | R:S | *-0.09 ± 0.02* | -0.143 | -0.034 | -3.204 | 0.002 |
| | Symbiosis M+ | -6.9 ± 2.1 | -11.125 | -2.652 | -3.185 | 0.002 |

Regression slopes are indicated in italics.

When it came to LAR, the negative relationship was independent of any variable, whereas in the case of RMF and R:S, it depended on the AM symbiosis, as the presence of AM fungi reduced in 16.1% and 6.9 % the magnitude of TDW variations for RMF and R:S ratio, respectively (Table 2). In the case of the RLR, this variable interacted with the presence of AM symbiosis in the plant, leading to a higher TDW percentage of change per unit of RLR plasticity, with respect to non-AM treatment, as revealed by the steeper slope (with 0.275 differential of coefficient estimate, Table 2 and Figure 3).

**Figure 3.**
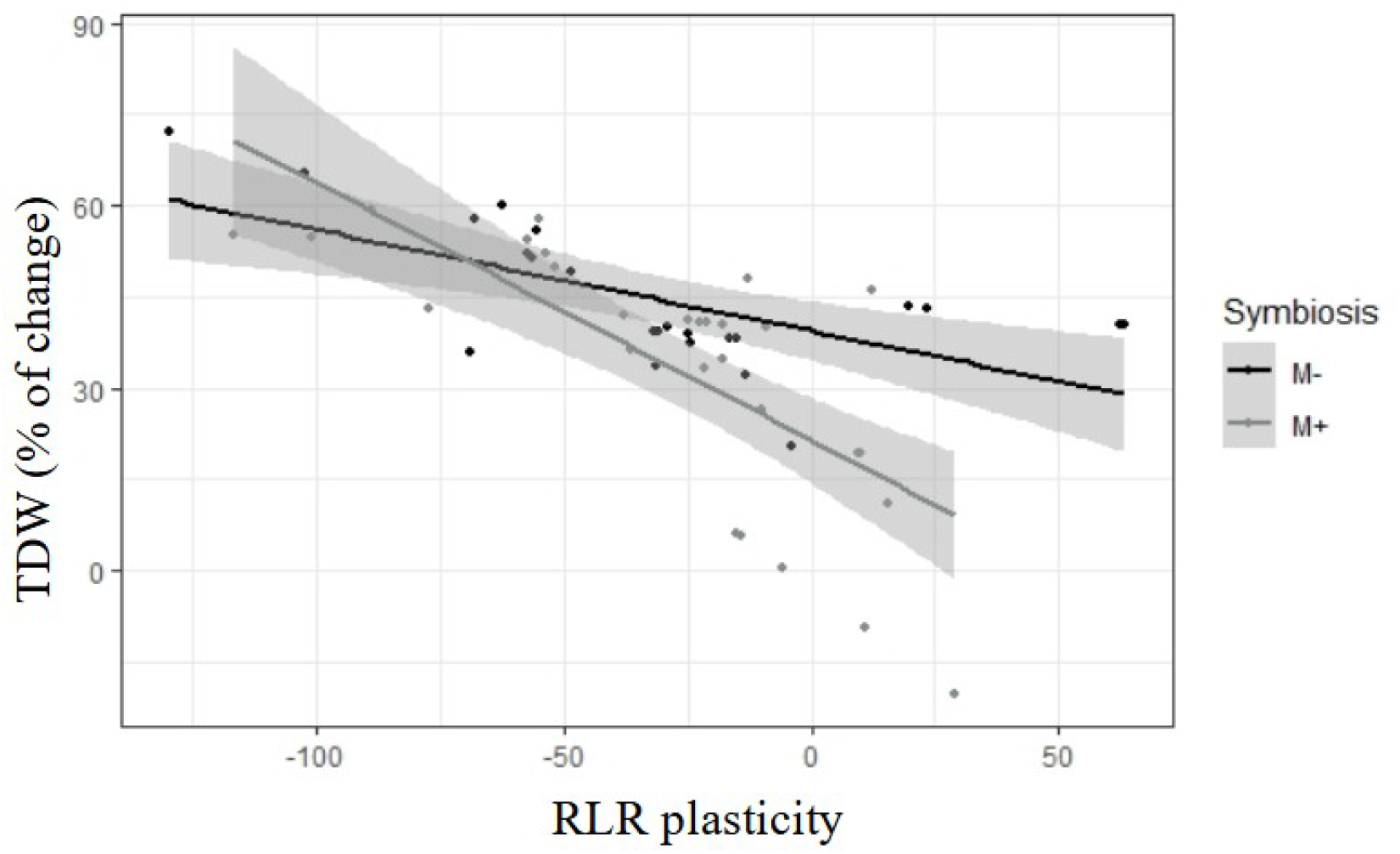
Linear mixed effects regression analysis of TDW percentage of change using RLR plasticity and AM symbiosis as independent variables. Data from salinity and drought stresses were pooled

## Discussion

The aim of the present work was to explore to what extent the presence of AM fungi alters the stress-induced plasticity of functional traits. Results from our meta-analysis revealed that the AM-symbiosis reduced the level of this plasticity in three out of seven below and aboveground plant functional traits retrieved from the literature. One of these functional traits was the LAR, which determines how much leaf area is present per unit of plant mass. Our meta-analysis result showing a lower magnitude of salt-induced LAR plasticity in AM plants, with respect to non-inoculated plants (Figure 1) agrees with findings from one field experiment performed with *Physalis peruviana* (cape gooseberry) growing on saline soil (Miranda et al., 2011). As far as we know, studies evaluating the influence of AM fungi on LAR plasticity of plants grown under soil-borne stress are limited to these two works. In contrast, there are comprehensive works that shed light on the effect of salinity over the SLA and LMF, the two LAR components. Response curves constructed from experiments under controlled conditions showed that increased environmental salinity reduces the SLA (leaves area/leaves mass; Poorter et al., 2009), whereas LMF (leaves mass/plant TDW) shows no variation with increased salinity (Poorter et al., 2012). This means that the overall effect of salinity across plant species is a reduction of the leaf area with respect to the plant biomass (LAR reduction). Taking this into consideration, for the salt-induced LAR plasticity to be negative (the most frequent case in the non-AM plants of our meta-analysis), the leaf area needed to be reduced by salinity to a lesser extent than plant biomass did. In contrast, AM plants were prone to reduce their leaf area to a higher extent than they reduced their TDW. One possible explanation for these results is that AM fungi have a beneficial effect on the leaf osmotic adjustment and CO_2_ assimilation (Augé 2004; Augé 2014; Evelin et al., 2019), processes generally hampered by salinity (Martinez-Ballesta et al., 2004; Heuer, 2005). This effect would release plants from investing more resources in leaf area to achieve carbon–nutrient colimitation, i.e.: to acquire carbon and nutrients maximizing plant benefits while minimizing resource acquisition costs (Maire et al., 2013). At a larger scale, as leaves are the seat of potential photosynthesizing and respiring plant components, the observed AM effect suggests that this symbiotic association might modulate energy balance (gains and expenditures) in agro-ecosystems from saline environments.

As our meta-analysis did not show significant effects by the AM symbiosis (nor by the type of stress) on SLA and LMF plasticities, we are impeded from further assessing the possible contribution of these components to the observed AM effect on LAR plasticity.

The other two functional traits whose stress-induced plasticities levels were afected by the AM-symbiosis, the R:S ratio and the RMF are conceptually similar as both reflect a different sensitivity of roots towards drought and salinity, compared to shoot (Munns and Tester 2008, Franco et al., 2011).

Our meta-analysis showed that, regardless the stress type, negative values of RMF and R:S ratio plasticity were the rule for non-AM plants (Figure 2), meaning that in those plants, the root biomass was reduced by the stress to a lesser extent than plant biomass and the shoot did (respectively). These results agree with comprehensive studies showing that in most plant species, RMF and R:S increase upon drought (Franco et al., 2011; Poorter et al., 2012; Eziz, 2017), and that salinity generally increases RMF and R:S ratio in glycophytes (Franco et al., 2011). In other words, the overall effect of drought and salinity across plant species is a lower reduction of the root biomass with respect to the shoot biomass (R:S ratio increase), or the plant biomass (RMF increase). The observed lower sensitivity of roots to these stresses would be a consequence of a rapid osmotic adjustment of roots, driven by increased biomass investment in the roots (Tang et al., 2022) and enhanced loosening ability of root cell walls (Sharp et al. 2004).

Our meta-analysis also revealed that plant species responded to the AM inoculation by reducing the magnitude of their stress-induced RMF and R:S ratio plasticities (less negative plasticity values), implying an improvement of shoot growth with respect to root growth performance under the stress, compared with non-AM plants. This result is in line with a previous study where the overall effect of AM colonization on the R:S ratio was analyzed (based on 11 trials; Veresoglou et al., 2012). We have also found that in some plant species, the improvement induced by AM inoculation exceeded a certain threshold, leading to positive values of RMF and R:S ratio plasticity. For the last observed AM effect on RMF and R:S ratio plasticities to occur, root biomass needs to be reduced, and/or shoot biomass increased by the stress to a higher extent, compared to non-AM plants. Decreases in the R:S ratio have been assigned to the alleviation of host nutrient limitation as a result of AM fungal establishment (e.g., Smith and Read 2008), which would reduce the need for root biomass investment. Such alleviation could be due to the fact that the extramatrical hyphae of AMF can increase the supply of water (Püschel et al., 2020) and ions (Marschner and Dell, 1994).

In parallel, despite changes in functional traits were weakly associated with variations in the total plant biomass (TDW), we observed that for a certain level of TDW reduction, AM plants responded with a lower RMF or R:S plasticity than non-AM ones (Table 2), contributing to idea that the AM symbiosis diminishes plant sensitivity to stress. The same observation can be made for the TDW/RLR regression but restricted to the -50% to 50% range of RLR plasticity, as the opposite effect was observed below -50%. This result is difficult to be interpreted considering the current knowledge and results obtained in the present meta-analysis. The RLR (which determines the root nutrient acquisition capacity) is composed by the SRL and the RMF (Eissenstat, 1997; Hill et al., 2006; Ostonen et al., 2007). Our meta-analysis could detect plasticity variations in the RMF, but not in the SRL component. It has been shown that under low nutrient conditions (as that attained by salinity and drought), the relative contribution of RMF to the increase of plant RLR, turns out to be far more important than the relative change in SRL (Freschet et al., 2015). However, more studies about the AM effect on stress-induced plasticity, specifically addressing SLA, LMF, SRL and RMF are needed for further analyzing their contribution to LAR and RLR. Ideally, these studies should contain data on biochemical parameters acknowledged as stress markers (such as proline), in order to analize their association with the observed changes in functional traits plasticity (insufficient in the present literature search). Interestingly, a wider database might allow the inclusion within the analysis of other predictor variables such as the AM fungal species, plant species, plant functional groups, and annual versus perennial, or wild versus domesticated plants.

### Agricultural implications

Most of the 71 species/cultivars included in the study are important food, industrial or forage crops (Supplementary Table 5). This proportion was expected as the study of AM symbiosis has generally been biassed by the agronomical viewpoint. Many of these economically important crops are often subjected to breeding practices directed to achieve yield stability over a range of environmental conditions, sometimes at the expense of reducing plasticity (Semchenko and Zobel 2005). However, the value of preserving plasticity during crop breeding has been progressively acknowledged (Matesanz and Milla, 2018; Sadras and Denison 2016).

Direction and magnitude of plant phenotypic changes in functional traits, as response to abiotic stresses are relevant information from the agro-ecological viewpoint. It has been stated that “yield should be higher if all individuals allocate less to competitive structures and functions than if all individuals respond to competition by allocating more resources to competitive structures (Weiner 2003; 2004)”. On this basis, it could be argued that the AM-induced lower proportion of resources invested in root than in shoot biomass as response to stress (less negative RMF plasticity, Fig. 1), is advantageous in crop production systems. Therefore, implementing agronomic practices that increase AM fungi propagules or promote the establishment of the AM symbiosis would lead to removing or reducing RMF plasticity, maximizing allocation to harvestable plant parts. In addition, the AM effects on plasticity detected in the present meta-analysis highlight the relevance of including AM fungi in programs aimed at selecting genotypes to be cultivated in the context of above-mentioned environmental constraints. Also, models designed to predict responses of specific crops to environmental conditions should incorporate changes induced by AM symbiosis on above versus belowground biomass allocation. The last would be also relevant from the edaphic viewpoint, as different plant organs vary in their decomposition rates: variations in functional traits may alter carbon trade-off between above and below ground, thus influencing soil organic matter build-up and nutrient recycling (Freschet et al., 2013).

Studies on the effects of fungal microsymbionts on plant phenotypic response induced by environmental restrictions evaluating changes in functional traits are limited and encompass few species or cultivars. For example, the impact of AM fungi on the root system plasticity (specific root length and proportion of fine roots) was studied on six maize varieties and found to constitute the most important adaptive strategy for maize to variation in P supply (Wang et al., 2020). Previously, endophytic fungi were shown to influence phenotypic plasticity responses of *Lolium perenne* to variable soil nutrients (Cheplick, 1997). Our meta-analysis spanning a significant number of species and cultivars, puts forward the notion of an overall modulating effect by AM symbiosis on plant plastic response to soil born abiotic stressors, although the different directions assumed by those plastic changes diverged across plant species for some functional traits. The divergence probably reflects species/cultivars widespread origins and adaptation strategies to the corresponding climates (Valladares and Sánchez-Gómez 2006), different plant phenological stages, or resources becoming restricted over the time lapse experiment (Poorter et al., 2012).

## Conclusions

In the present meta-analysis, we quantified the size effect of AM symbiosis on the stress-induced plasticity of several reported and calculated functional traits, using linear mixed model analysis. Our results provided evidence to accept the hypothesis that AMF mycorrhizal inoculation may reduce the phenotypic plasticity of important plant functional traits (LAR, RMF and R:S ratio), by decreasing its magnitude. We also found a weak correlation between traits plasticity and total biomass variation. Although we believe our literature search and data collection were intensive and our results robust, the scope of our conclusions is limited by the agronomical bias of plant species targeted by the meta-analysis. Further knowledge on non-cultivable plant species and better understanding of the mechanisms ruling resources allocation in plants would allow more generalized conclusions.

## Author’s Contributions

Ana Menéndez and David Bilenca contributed to the study conception and design. Florencia Gobbo, César Bordenave and Ayelén Gázquez contributed with data collection; statistical analysis was performed by María José Corriale. The first draft of the manuscript was written by Ana Menéndez and all authors commented on previous versions of the manuscript. All authors read and approved the manuscript

**Supplementary Table 1.**
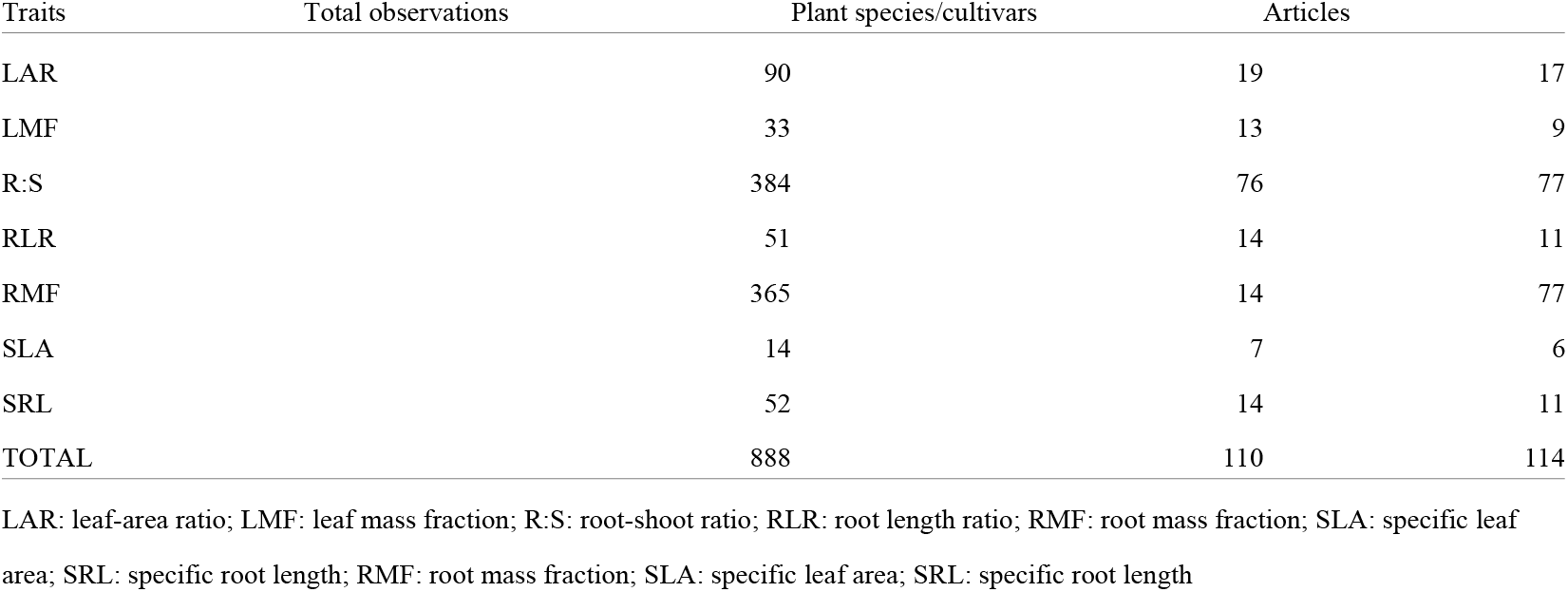
Summary of the number of observations, plant species and articles where the trait was extracted.

**Supplementary Table 2.** Candidate models accounting for variations in functional traits plasticities. Selected model are shown in bold

|  |  |  |  |  |  |  |
| --- | --- | --- | --- | --- | --- | --- |
| Functional traits | Candidate models | Predictors | K | AIC | Wald's test |  |
| RMF | RMF1 | Estrés*Symbiosis | 9 | 3429.3 |  |  |
| | RMF2 | Estres+Symbiosis | 7 | 3428.2 | $\chi^2$ 1-2=2.84; $p=0.24$ | |
|  | RMF3 | Symbiosis | 5 | 3429.0 | **$\chi^2$ 2-3=2.84; $p=0.09$** |  |
| | RMF4 | null | 4 | 3431.9 | $\chi^2$ 3-4=4.96; $p=0.02$ | |
| LAR | LAR1 | Estrés*Symbiosis | 7 | 863.9 |  |  |
| | LAR2 | Estres+Symbiosis | 6 | 880.8 | $\chi^2$ 1-2=18.01;<br>$p<0.001$ | |
| LMF | LMF1 | Estrés*Symbiosis | 7 | 287.9 |  |  |
| | LMF2 | Estres+Symbiosis | 6 | 286.5 | $\chi^2$ 1-2=0.58; $p=0.44$ | |
| | LMF3 | Symbiosis | 5 | 286.8 | $\chi^2$ 2-3=2.33; $p=0.12$ | |
| | LMF4 | null | 4 | 284.8 | $\chi^2$ 3-4=0.005; $p=0.94$ | |
| SRL | SRL1 | Estrés*Symbiosis | 7 | 607.7 |  |  |
| | SRL2 | Estres+Symbiosis | 6 | 607.1 | $\chi^2$ 1-2=1.35; $p=0.24$ | |
| | SRL3 | Symbiosis | 5 | 605.6 | $\chi^2$ 2-3=0.56; $p=0.45$ | |
| | SRL4 | null | 4 | 604.5 | $\chi^2$ 3-4=0.81; $p=0.36$ | |
| RLR | RLR1 | Estrés*Symbiosis | 7 | 513.4 |  |  |
| | RLR2 | Estres+Symbiosis | 6 | 514.9 | $\chi^2$ 1-2=3.45; $p=0.06$ | |
| | RLR3 | Symbiosis | 5 | 512.9 | $\chi^2$ 2-3=0.07; $p=0.78$ | |
| | RLR4 | null | 4 | 514.0 | $\chi^2$ 3-4=0.81; $p=0.36$ | |
| S:R | S:R1 | Estrés*Symbiosis | 9 | 3954.5 |  |  |
| | S:R2 | Estres+Symbiosis | 7 | 3959.1 | $\chi^2$ 1-2=4.25; $p=0.06$ | |
|  | S:R3 | Symbiosis | 5 | **3963.8** | **$\chi^2$ 2-3=3.11; $p=0.07$** |  |
| | S:R4 | null | 4 | 3987.4 | $\chi^2$ 3-4=25.64;<br>$p<0.001$ | |
| SLA | SLA1 | Estrés*Symbiosis | 7 | 92.6 |  |  |
| | SLA2 | Estres+Symbiosis | 6 | 96.4 | $\chi^2$ 1-2=0.17; $p=0.67$ | |
| | SLA3 | Symbiosis | 5 | 102.0 | $\chi^2$ 2-3=0.51; $p=0.47$ | |
| | SLA4 | null | 4 | 103.8 | $\chi^2$ 3-4=0.11; $p=0.73$ | |
| K: number of parameters; AIC: Akaike's Information Criterion and Wald's tatistic values. The best model (with the lowest AIC value) is shown in bold. |  |  |  |  |  |  |

**Supplementary Table 3.** Percentage of variance explained by random effects.

| Trait | Variance (%) |  |  |
| --- | --- | --- | --- |
|  | Plant species/cultivars | Articles | Residue |
| LAR | 1 | 38 | 61 |
| LMF | 0 | 54 | 46 |
| RLR | 0 | 67 | 33 |
| RMF | 0 | 44 | 56 |
| R:S | 0 | 29 | 71 |
| SLA | 4 | 88 | 9 |
| SRL | 0 | 76 | 24 |

LAR: leaf area ratio; LMF: leaf mass fraction; RLR: root length ratio;
RMF: root mass fraction; R:S: root-shoot ratio; SLA: specific leaf area; SRL: specific root length

**Supplementary Table 4.** Candidate models for associations between TDW variation and plasticity of functional traits. Data from salinity and drought stresses were pooled.

| Functional traits | Candidate models | Predictors | K | AIC | Wald's test |
| --- | --- | --- | --- | --- | --- |
| LAR | LAR1 | LAR*Symbiosis |  | 6784.5 |  |
| | LAR2 | LAR+Symbiosis | | 5783.1 | $\chi^2$ 1–2=0.49; $p=0.48$ |
|  | LAR3 | LAR |  | **4783.3** | **$\chi^2$ 2–3=2.32; $p=0.12$** |
| | LAR4 | null | | 3814.6 | $\chi^2$ 3–4=25.78; $p<0.001$ |
| RLR | RLR1 | RLR*Symbiosis |  | 7863.9 |  |
| | RLR2 | RLR+Symbiosis | | 6880.8 | $\chi^2$ 1–2=18.01; $p<0.001$ |
| RMF | RMF1 | RMF*Symbiosis |  | 63120.2 |  |
|  | RMF2 | RMF+Symbiosis |  | **53118.2** | **$\chi^2$ 1–2=0.08; $p=0.77$** |
| | RMF3 | RMF | | 43125.5 | $\chi^2$ 2–3=9.28; $p=0.002$ |
| R:S | RS1 | RS*Symbiosis |  | 72961.8 |  |
|  | RS2 | RS+Symbiosis |  | **62960.0** | **$\chi^2$ 1–2=0.27; $p=0.270$** |
| | RS3 | Symbiosis | | 52960.0 | $\chi^2$ 2–3=10.04; $p=0.002$ |

LAR: leaf area ratio; RLR: root length ratio; RMF: root mass fraction; R:S: root-shoot ratio; K: number of
parameters; AIC: Akaike's Information Criterion and Wald's statistic values. The best model (with the lowest AIC value and lowest $p$ ) is shown in bold.

